# Transcriptomic analysis of dystonia-associated genes reveals functional convergence within specific cell types and shared neurobiology with psychiatric disorders

**DOI:** 10.1101/2020.01.31.928978

**Authors:** Niccolò E. Mencacci, Regina Reynolds, Sonia Garcia Ruiz, Jana Vandrovcova, Paola Forabosco, UK Brain Expression Consortium, International Parkinson’s Disease Genomics Consortium, Michael E. Weale, Kailash P. Bhatia, John Hardy, Juan A Botía, Mina Ryten

## Abstract

Dystonia is a neurological disorder characterized by sustained or intermittent muscle contractions causing abnormal movements and postures, often occurring in absence of any structural brain abnormality. Psychiatric comorbidities, including anxiety, depression, obsessive-compulsive disorder and schizophrenia, are frequent in dystonia patients. While mutations in a fast-growing number of genes have been linked to Mendelian forms of dystonia, the cellular, anatomical, and molecular basis remains unknown for most genetic forms of dystonia, as does its genetic and biological relationship to neuropsychiatric disorders. Here we applied an unbiased systems-biology approach to explore the cellular specificity of all currently known dystonia-associated genes, predict their functional relationships, and test whether dystonia and neuropsychiatric disorders share a genetic relationship. To determine the cellular specificity of dystonia-associated genes in the brain, single-nuclear transcriptomic data derived from mouse brain was used together with expression-weighted cell-type enrichment. To identify functional relationships amongst dystonia-associated genes, we determined the enrichment of these genes in co-expression networks constructed from ten human brain regions. Stratified linkage-disequilibrium score regression was used to test whether co-expression modules enriched for dystonia-associated genes significantly contribute to the heritability of anxiety, major depressive disorder, obsessive-compulsive disorder, schizophrenia, and Parkinson’s disease. Dystonia-associated genes were significantly enriched in adult nigral dopaminergic neurons and striatal medium spiny neurons. Furthermore, four of the 220 gene co-expression modules tested were significantly enriched for the dystonia-associated genes. The identified modules were derived from the substantia nigra, putamen, frontal cortex, and white matter, and were all significantly enriched for genes associated with synaptic function. Finally, we demonstrated significant enrichments of the heritability of depression, obsessive-compulsive disorder and schizophrenia, but not anxiety and Parkinson’s disease, within the putamen and white matter modules. In conclusion, multiple dystonia-associated genes interact and contribute to pathogenesis likely through dysregulation of synaptic signalling in striatal medium spiny neurons, adult nigral dopaminergic neurons and frontal cortical neurons. Furthermore, the enrichment of the heritability of psychiatric disorders in the co-expression modules enriched for dystonia-associated genes indicates that psychiatric symptoms associated with dystonia are likely to be intrinsic to its pathophysiology.

## Introduction

The term dystonia defines a heterogeneous family of hyperkinetic movement disorders unified by their clinical manifestations, which include sustained or intermittent muscle contractions causing abnormal, often repetitive, movements, postures, or both (Albanese *et al*., 2013). While dystonia can occur as a consequence of both focal and degenerative brain lesions, most commonly affecting the basal ganglia (Bhatia and Marsden, 1994) or cerebellum (Batla *et al*., 2015), the majority of patients with dystonia have normal neuroimaging findings and pathological studies have consistently shown the absence of even subtle structural abnormalities (Sharma, 2019). Thus, like epilepsy, dystonia has a dual nature, presenting both as the symptom of specific brain lesions and as a discrete disease entity.

The etiology of dystonia occurring in the absence of structural abnormalities is unknown in most cases. Pathogenic mutations in a fast-growing number of genes have been linked to Mendelian forms of “idiopathic dystonia” and appear to be an important cause of dystonia, especially in cases with pediatric onset of symptoms and/or strong family history (Balint *et al*., 2018). Clinically, mutations in these dystonia-associated genes (from hereafter termed DYT genes) produce a range of phenotypes, including isolated (dystonia occurring alone), combined (dystonia associated with myoclonus or parkinsonism) and paroxysmal forms of the condition (intermittent attacks of dystonia with normal interictal neurological examination) (Marras *et al*., 2016; Lohmann and Klein, 2017; Zhang *et al*., 2019), with a subset of genes capable of producing multiple clinical presentations (Friedman *et al*., 2016; Carecchio *et al*., 2017; Zech *et al*., 2017; Balint *et al*., 2019).

Genetic discoveries played a key role in conclusively pushing dystonia into the realm of neurological disorders after decades of controversy during which dystonia was considered by many to be a psychiatric condition (Lesser and Fahn, 1978). Still, dystonia not only affects motor function, but presents with additional psychiatric symptoms in 50-90% of patients with dystonia, including both patients with sporadic and monogenic forms (Stamelou *et al*., 2012; Peall *et al*., 2013; Zurowski *et al*., 2013; Peall *et al*., 2015; Conte *et al*., 2016). These symptoms range from depression and anxiety (Lencer *et al*., 2009; Fabbrini *et al*., 2010; Steinlechner *et al*., 2017) to obsessive compulsive disorder (Voon *et al*., 2010; Barahona-Correa *et al*., 2011) and psychosis (Brashear *et al*., 2012; Vijiaratnam *et al*., 2018; Timmers *et al*., 2019), with depression and anxiety disorders being the most common (Smit *et al*., 2016; Berman *et al*., 2017). Psychiatric co-morbidities often precede the onset of motor symptoms (Moraru *et al*., 2002; Lencer *et al*., 2009) and have a profound impact on dystonia patients’ quality of life (Smit *et al*., 2016; Eggink *et al*., 2019).

These observations raise the question of whether psychiatric symptoms are intrinsic to the neurobiology of dystonia, a question that is hard to address as the physiological and pathogenic roles of DYT genes are known only for a minority of genes. For instance, it is well established that the genes associated with forms of dystonia responsive to dopamine replacement (i.e. DOPA-responsive dystonias) are all involved in the synthesis and metabolism of dopamine in nigral dopaminergic neurons (Ng *et al*., 2015; Ribot *et al*., 2019). However, little is known about the biological function of many of the other DYT genes and how they contribute to disease pathogenesis.

The recent progress in our understanding of the genetic architecture of both dystonia and neuropsychiatric diseases, together with the increased availability of brain-related functional genomic annotations, offers a unique opportunity to robustly examine the relationship between these conditions. More specifically, whole-exome sequencing efforts have resulted in the identification of several novel monogenic causes of dystonia (described above), and genome wide association studies (GWAS) for schizophrenia (Pardinas *et al*., 2018), obsessive-compulsive disorder (IOCDF-GC and OCGAS, 2018), anxiety (Otowa *et al*., 2016), and major depressive disorder (Wray *et al*., 2018) have provided an increasingly long list of risk loci.

Thus, in this study we applied a comprehensive systems biology approach to explore the following unanswered questions: (i) Which brain cells are most relevant to the pathogenesis of monogenic dystonias? (ii) Do DYT genes interact and coalesce in shared cellular and molecular pathways? And if yes, in which brain regions and/or cells does this happen? (iii) Is there a genetic relationship between dystonia and neuropsychiatric disorders suggesting a shared neurobiological basis? Setting aside the DYT genes contributing to forms of DOPA-responsive dystonia, our analyses suggest that multiple dystonia-genes interact and play a fundamental role in the pathogenesis of dystonia likely through dysregulation of synaptic signalling in striatal medium spiny neurons (MSNs) and frontal cortex pyramidal neurons. Furthermore, we show that DYT genes enrich within co-expression modules that also enrich for the heritability of neuropsychiatric disorders, suggesting that the psychiatric symptoms associated with dystonia are intrinsic to its pathophysiology.

## Methods

### Definition of a list of “idiopathic dystonia”-related genes

The Movement Disorders Society (MDS) has recently provided a list of the established genes clinically associated with prominent dystonia (Marras *et al*., 2016). From this list, we included only those genes that when mutated cause prominent dystonia in the absence of imaging or neuropathological evidence of structural or degenerative abnormalities (as reported by at least two independent groups). For this reason, all DYT genes systematically associated with a radiological or pathological phenotype of basal ganglia injury (i.e. mitochondrial, lysosomal storage, or metabolic disorders), damage due to metal accumulation (i.e. iron, manganese, copper and calcium) or overt neurodegeneration (e.g. *TAF1* and other genes associated with degenerative complex dystonia-parkinsonism) were excluded. Furthermore, we included all DYT genes regardless of their most commonly associated clinical phenotype (i.e. isolated, combined and paroxysmal dystonia). To ensure our list of DYT genes was up to date, we complemented the MDS list with the additional confirmed isolated and combined dystonia genes (*ANO3* and *KMT2B*) included in the most recently published review on dystonia (Balint *et al*., 2018). Furthermore, we included four additional genes for which a established pathogenic role in dystonia has only been recently demonstrated, namely *KCTD17* (Mencacci *et al*., 2015; Graziola *et al*., 2019; Marce-Grau *et al*., 2019), *HPCA* (Charlesworth *et al*., 2015; Atasu *et al*., 2018), *SLC18A2* (Rilstone *et al*., 2013; Rath *et al*., 2017), *DNAJC12* (Anikster *et al*., 2017; Veenma *et al*., 2018). As for paroxysmal dystonias, we included the confirmed genes listed in the most recent review on the topic (Zhang *et al*., 2019). The confirmed DYT genes included in the analysis are listed in Table 1. Unconfirmed DYT genes (i.e. *CIZ1*, *COL6A3, RELN*) were not included in the analysis.

**Table 1.**
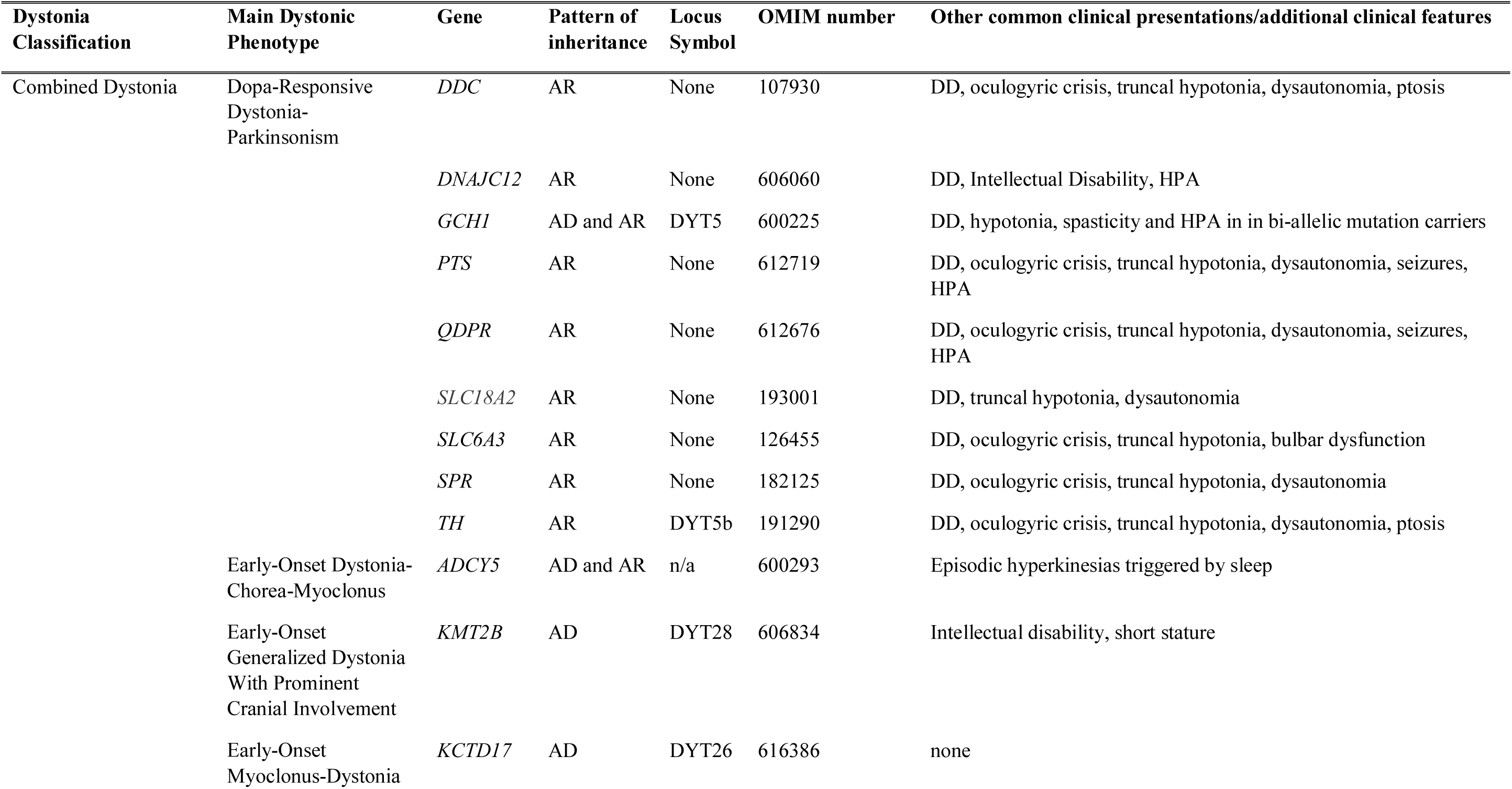
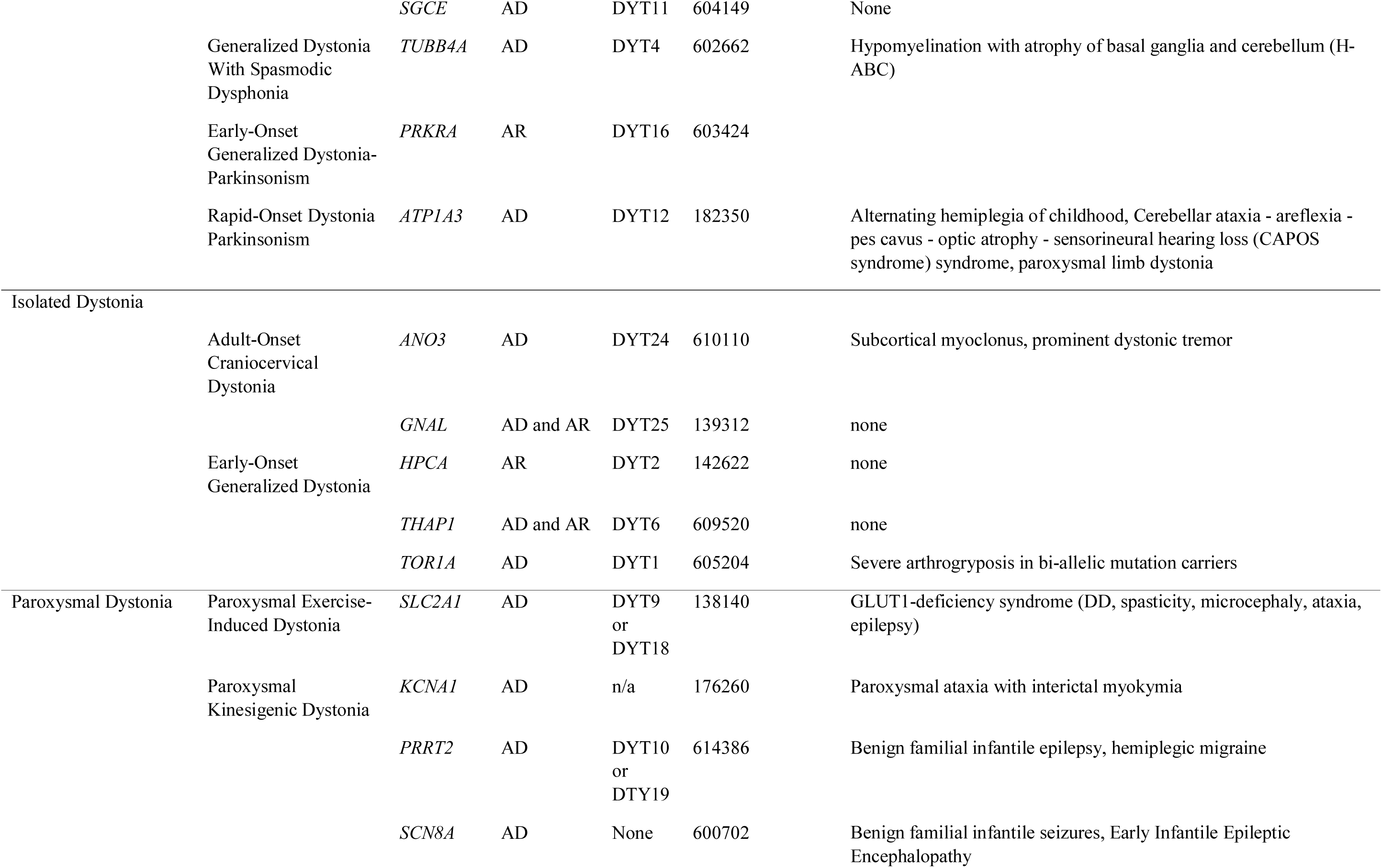
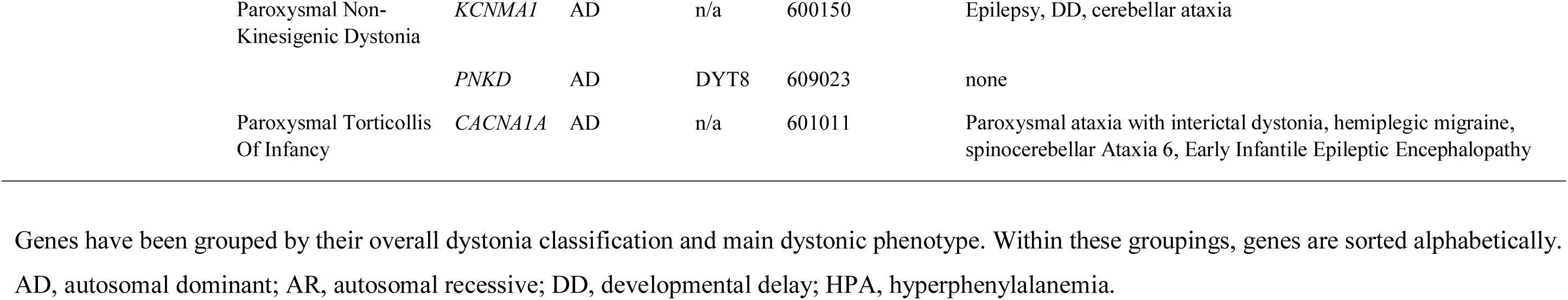
List of dystonia-associated genes included in the analysis.

### Expression-weighted cell-type enrichment (EWCE)

EWCE (see URLs) was used to determine whether DYT genes have higher expression within particular brain-related cell types than would be expected by chance (Skene and Grant, 2016). As our input we used 1) the list of DYT genes as defined above and 2) specificity values calculated by Skene et al for level 1 cell types from the Karolinska single-cell RNA-sequencing superset, which includes cell types from the neocortex, hippocampus, hypothalamus, striatum and midbrain (see URLs) (Skene *et al*., 2018). EWCE with the target list was run with 100,000 bootstrap replicates, which were sampled from a background list of genes that excluded all genes without a 1:1 mouse:human ortholog. We additionally controlled for transcript length and GC-content biases by selecting bootstrap lists with comparable properties to the target list. Data are displayed as standard deviations from the mean, and any values < 0, which reflect a depletion of expression, are displayed as 0. P-values were corrected for multiple testing using the Benjamini-Hochberg method over all cell types.

### Weighted Gene Co-Expression Analysis (WGCNA)

We generated Gene Co-expression Networks (GCN) for CNS tissue-specific transcriptomic data generated by the UK Brain Expression Consortium (Forabosco *et al*., 2013) and Genotype-Tissue Expression Consortium (2015) (Version 6; www.gtexportal.org). In total, 57 gene-level expression datasets across an equal number of tissues were used with the weighted gene co-expression network analysis (WGCNA) R package with k-means adjustment to generate tissue-specific networks (Langfelder and Horvath, 2008; Langfelder *et al*., 2008; Botia *et al*., 2017). For each tissue, a “signed” GCN was constructed by creating a signed Topological Overlap Measure (TOM) matrix based on Pearson correlation. Gene modules were created by hierarchical clustering based on a 1-TOM dissimilarity matrix. The results of the initial hierarchical clustering were post-processed using the k-means clustering search method with 30 iterations. All co-expression networks are available online via the CoExp website (https://snca.atica.um.es/coexp/Run/Catalog/) or CoExpNets package (https://github.com/juanbot/CoExpNets) to enable use with third-party software.

To assess module preservation within and between datasets we performed a preservation analysis based on WGCNA using the Z summary statistic to evaluate preservation (Langfelder *et al*., 2011). As per Langfelder *et al*., if Z summary statistic > 10, there is strong evidence of module preservation; if 2 < Z summary statistic < 10, there is weak to moderate evidence of preservations; and finally, if Z summary statistic < 2, there is no evidence of preservation.

Gene modules were functionally annotated with gProfileR R package using Gene Ontology (GO) database without Electronic Inferred Annotations (EIA) and accounting for multiple testing with gSCS (Reimand *et al*., 2007). Additional functional annotations for modules of interest were generated using the web server SynGO (https://www.syngoportal.org/), which provides an expert-curated resource for synapse function and gene enrichment analysis (Koopmans *et al*., 2019). All gene set enrichment analyses were performed using default settings, with no annotation filters applied and a minimum gene count of three for ontology terms to be included in the overrepresentation analysis.

### Stratified linkage disequilibrium score regression (LDSC)

Stratified LDSC (see URLs) (Finucane *et al*., 2015) was used to test whether co-expression modules enriched for DYT genes significantly contributed to the common SNP heritability of four neuropsychiatric disorders (anxiety, major depressive disorder; obsessive compulsive disorder; and schizophrenia) and one neurodegenerative disorder (Parkinson’s disease) (Table 2). Co-expression modules from UKBEC were downloaded using the CoExpNets package (see URLs). As previously performed, modules were filtered to include only genes with module membership ≥ 0.5 (Reynolds *et al*., 2019). Furthermore, gene coordinates were extended by 100kb upstream and downstream of their transcription start and end site, in order to capture regulatory elements that might contribute to disease heritability (Finucane *et al*., 2018).

**Table 2.** Summary of GWAS datasets.

| **Disease** | **First author, Year** | **N cases** | **N controls** | **Reference** |
| --- | --- | --- | --- | --- |
| Anxiety | Otowa, 2016 | 7,016 | 14,745 | (Otowa *et al.*, 2016) |
| Major depressive disorder (excluding 23andMe contributions) | Wray, 2018 | 59,851 | 113,154 | (Wray *et al.*, 2018) |
| Obsessive compulsive disorder | IOCDF-GC, 2017 | 2,688 | 7,037 | (IOCDF-GC and OCGAS, 2018) |
| Parkinson's disease (excluding 23andMe contributions) | Nalls, 2019 | 33,674<br>(18,618 proxy cases from UK Biobank) | 449,037 | (Nalls *et al.*, 2019) |
| Schizophrenia | Pardiñas, 2018 | 40,675 | 64,643 | (Pardinas *et al.*, 2018) |

All annotations were constructed in a binary format (1 if the SNP was present within the annotation and 0 if not), using all SNPs with a minor allele frequency > 5%. Annotations were then added individually to the baseline model of 53 annotations provided by Finucane et al. (version 1.1, see URLs), comprising genome-wide annotations reflecting genetic architecture. HapMap Project Phase 3 (HapMap3) (Altshuler *et al*., 2010) SNPs and 1000 Genomes Project (Abecasis *et al*., 2012) Phase 3 European population SNPs were used for the regression and LD reference panels, respectively. The MHC region was excluded from all analyses due to the complex and long-range LD patterns in this region. For all stratified LDSC analyses, we report a one-tailed p-value (coefficient p-value) based on the coefficient z-score outputted by stratified LDSC. A one-tailed test was used as we were only interested in annotation categories with a significantly positive contribution to trait heritability, conditional upon the baseline model. The significance threshold was set to 0.05 divided by the number of modules for each co-expression network (four for UKBEC) and the number of GWAS runs.

## Results

### Dystonia genes are highly and specifically expressed in midbrain dopaminergic and striatal medium spiny neurons

In the first instance, we wanted to study the cellular specificity of DYT genes, with the aim of identifying the cell types most likely involved in primary pathology. Using EWCE analysis and single-cell gene expression profiling of the mouse brain (Karolinska Institute brain superset from the Linnarsson group), we demonstrated that DYT genes are significantly enriched in two cell types, namely adult nigral dopaminergic neurons (FDR-adjusted p-value < 0.00001) and striatal MSNs (FDR-adjusted p-value = 0.00156; Fig. 1A; **Supplementary Table 1**).

**Figure 1.**
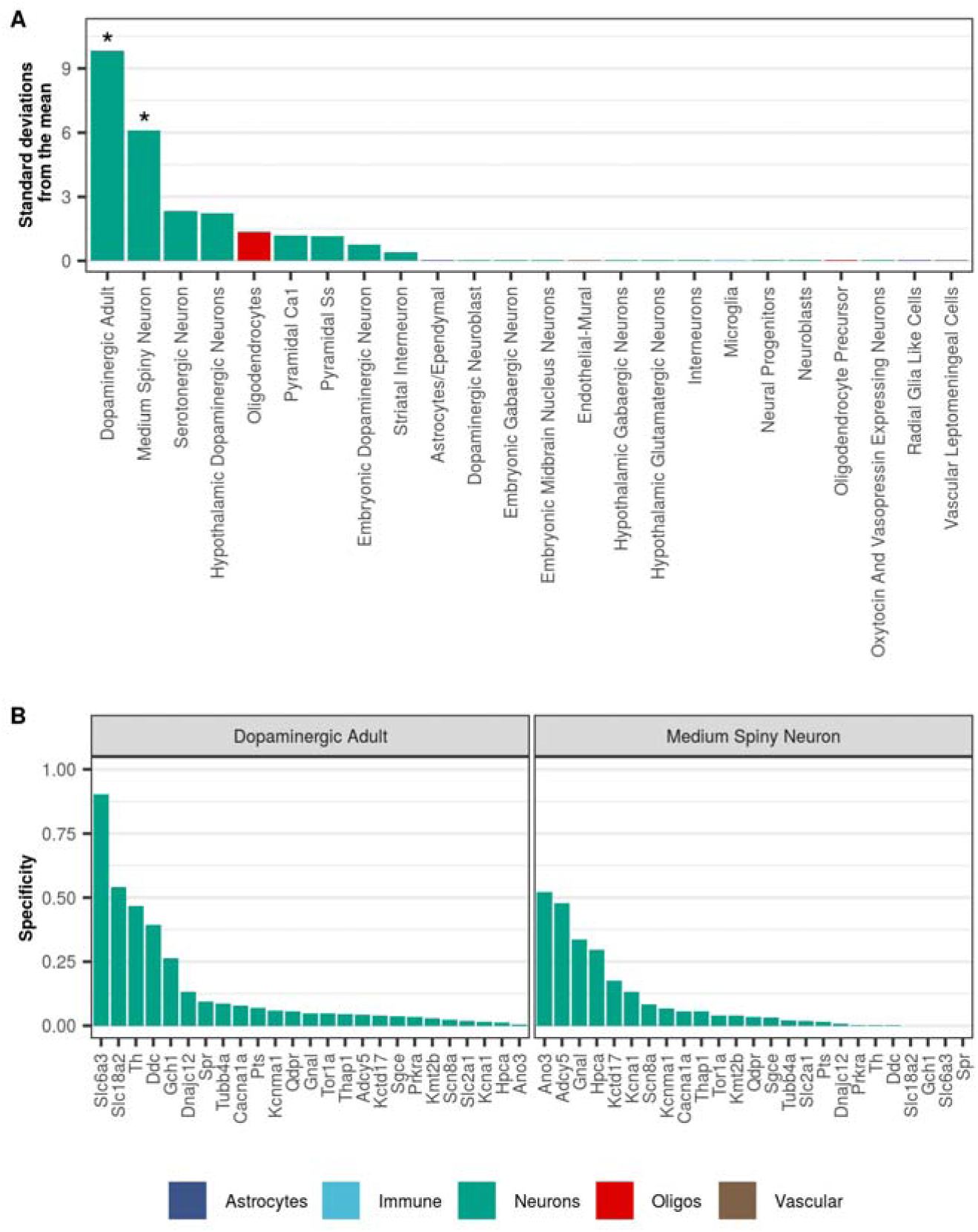
Dystonia-associated genes are highly expressed in dopaminergic neurons and medium spiny neurons. (**A**) Enrichment of dystonia-associated genes in level 1 cell types from the Karolinska superset was determined using EWCE. Standard deviations from the mean indicate the distance of the mean expression of the target list from the mean expression of the bootstrap replicates. Asterisks denote significance at p < 0.05 after correcting for multiple testing with the Benjamini-Hochberg method over all level 1 cell types. Numerical results are reported in **Supplementary Table 1**. (**B**) Plot of specificity values for all dystonia-associated genes within adult nigral dopaminergic neurons and medium spiny neurons (level 1 cell types from the Karolinska single-cell RNA-sequencing superset). Specificity values were derived from Skene et al. (Skene *et al*., 2018) who calculated specificity by dividing the mean expression of a gene in one cell type by the mean expression in all cell types. In other words, specificity is the proportion of a gene’s total expression attributable to one cell type, with a value of 0 meaning a gene is not expressed in that cell type and a value of 1 meaning that a gene is only expressed in that cell type. In both plots, cell types are coloured by the overall class they belong to (e.g. astrocyte, neuron, oligodendrocyte, etc.).

The enrichment within adult nigral dopaminergic neurons was largely driven by genes responsible for forms of dystonia responsive to dopaminergic therapies (i.e. DOPA-responsive dystonias) (Ng *et al*., 2015). In fact, we noted that *SLC6A3, SLC18A2, TH, DDC,* and *GCH1,* had the highest specificity values in adult nigral dopaminergic neurons (Fig. 1B**; Supplementary Table 1**).

*ANO3, ADCY5, GNAL,* and *KCTD17* were the genes with the highest specificity values in striatal MSNs. Amongst these, *ADCY5* and *GNAL* had specificity values more than 4-fold higher in MSNs than in other cell-types, indicating almost exclusive expression in this cell type (**Supplementary Fig. 1; Supplementary Table 1**). While some of the genes driving this enrichment have an established function in MSNs (i.e. *GNAL* and *ADCY5* are involved in striatal dopaminergic and adenosinergic post-receptor signaling) (Goodchild *et al*., 2013), *ANO3* and *KCTD17* have not been studied in this context and their physiological role in MSNs is currently unknown.

### Gene co-expression network analysis identifies dystonia-enriched modules

Commonalities in the cellular specificity of DYT genes suggested the possibility that a subset of DYT genes may be functionally related. However, EWCE analysis was based on cell-specific gene expression data derived from mouse and did not consider correlations in gene expression. Co-expression networks have proven to be an efficient means of identifying hidden functional relationships between genes of interest (Oldham *et al*., 2008; Kelley *et al*., 2018). Thus, to further explore the possibility that a subset of DYT genes are functionally related, we tested for the enrichment of DYT genes within co-expression networks generated from ten regions of the human brain (cerebellar cortex, frontal cortex, hippocampus, inferior olivary nucleus, occipital cortex, putamen, substantia nigra, temporal cortex, thalamus and intralobular white matter) using transcriptomic data provided by the UK Brain Expression Consortium (UKBEC) (www.rytenlab.com/coexp/Run/Catalog/) (Ramasamy *et al*., 2014).

Amongst the 220 gene co-expression modules tested, we identified four modules in which DYT genes were significantly enriched (FDR-adjusted p-value < 0.05; **Supplementary Table 2**). These four modules derived from the substantia nigra (“cyan” module), putamen (“cyan” module), frontal cortex (“lightyellow” module) and white matter (“blue” module) networks. No enrichment of DYT genes was observed in any of the other brain regions. To replicate these findings, we tested for enrichment of DYT genes in co-expression networks generated using transcriptomic data from the Genotype Tissue Expression Project (GTEx); only co-expression networks derived from frontal cortex, putamen and substantia nigra were used. Amongst the co-expression modules tested, DYT genes enriched in three modules: frontal cortex “turquoise” module, putamen “blue” module and substantia nigra “darkorange2” module.

The overlap of DYT genes with the substantia nigra “cyan” module was driven by seven DYT genes (*GCH1, TH, SLC6A3, DNAJC12, DDC, SLC18A2, PTS;* FDR-adjusted p-value = 0.0409) (Fig. 2A). This finding was replicated using GTEx-derived co-expression networks, with a significant enrichment of DYT genes in the substantia nigra “darkorange2” module (FDR-adjusted p-value = 0.005) driven by an overlapping set of six genes, namely *GCH1, TH, SLC6A3, DDC, SLC18A2* and *PTS*. Importantly, mutations in all DYT genes enriched in these modules cause DOPA-responsive dystonias and are well known to be functionally related to dopamine synthesis and/or metabolism.

**Figure 2.**
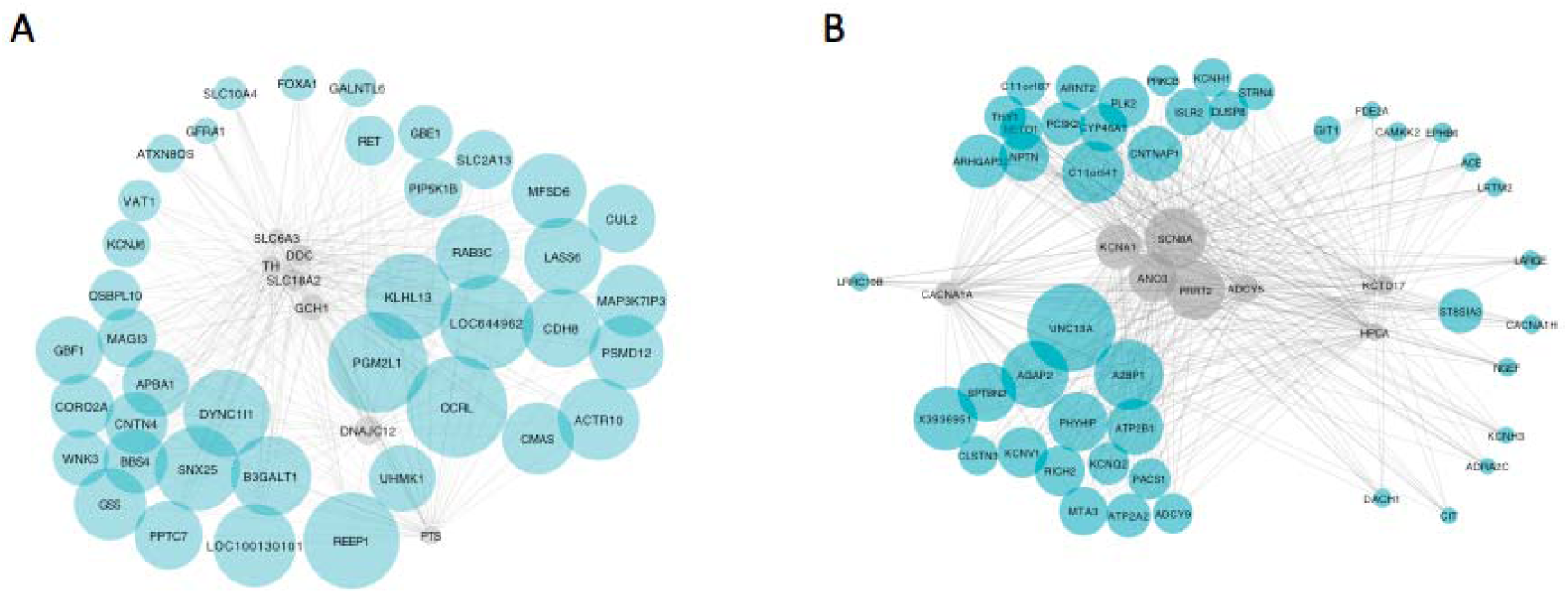
Gene co-expression modules enriched for dystonia genes in the substantia nigra and putamen. The substantia nigra “cyan” (**A**) and putamen “cyan (**B**) dystonia-linked UKBEC modules visualised using “bottom-up” plots. In each case, the dystonia genes within the modules are depicted as grey nodes with their most connected genes, as determined by the Topology Overlap Measure, depicted in cyan (to a maximum of 7 genes per seed). Node size reflects connectivity. Plots were generated using Cytoscape 3.5.1 and the Edge-weighted Spring Embedded layout algorithm was used for rendering to a 2D canvas.

The most significant enrichment of DYT genes was detected in a putamen gene co-expression module. Of the 28 DYT genes tested, eight overlapped with the “cyan” module in this tissue (*ADCY5, ANO3, KCTD17, HPCA, PRRT2, SCN8A, KCNA1, CACNA1A;* FDR-adjusted p-value = 0.0001) (Fig. 2B). This module included established cellular markers of MSNs, including *DRD1*, *DRD2*, *ADORA2A* and *PPP1R1B*, indicating the module captures the expression signature of these neurons. Again, this finding was supported by replication in GTEx-derived co-expression networks (putamen “blue” module; FDR-adjusted p-value = 0.026), with the enrichment driven by an overlapping set of six DYT genes (*ADCY5, KCTD17, HPCA, PRRT2,* and *CACNA1A*).

Significant enrichment of DYT genes was also detected in the white matter “blue” module (overlap of eight genes, *PRRT2, PNKD, SCN8A, KCNA1, ATP1A3, ANO3, KCTD17, HPCA*; FDR-adjusted p-value = 0.0203) and the “light-yellow” frontal cortex module (overlap of six genes*, ANO3, GNAL, KCTD17, HPCA, KCNMA1, CACNA1A;* FDR-adjusted p-value = 0.0203). Both modules were predicted to be expression signatures of cortical pyramidal neurons. Where analysis was possible due to data availability (white matter tissue is not available in GTEx), we noted that once again the findings in frontal cortex were replicated in GTEx-derived co-expression networks (frontal cortex “turquoise” module; p-value =0.0289).

Functional annotation showed that all four UKBEC dystonia-linked modules were significantly enriched for genes associated with neuronal synaptic transmission (putamen “cyan” module, corrected p-value = 3.44 x10^-8^; white matter “blue” module p-value = 3.4 x 10^-52^; frontal cortex “lightyellow” module, p-value = 9.82 x 10^-7^; substantia nigra cyan module, p-value = 0.0247). Furthermore, in the case of both the “blue” and “lightyellow” modules we found significant enrichment of terms relating to neurodevelopment, namely neuron projection development (white matter “blue” module, p-value = 4.96 x 10^-26^) and nervous system development (frontal cortex “lightyellow” module, nervous system development, p-value = 1.14 x 10^-5^). Finally, in case of the “cyan” putamen module, the most over-represented terms related to metal ion transmembrane transport, indicating an enrichment of genes coding for ion channels.

To investigate the significance of these modules further, we sought to identify whether the enrichment of genes relating to synaptic transmission was driven by genes involved in post-synaptic as compared to pre-synaptic structures. To address this question, we checked for enrichment of genes that have been reliably associated with a specific synaptic structure using the recently released SynGO database (Koopmans *et al*., 2019). This demonstrated significant enrichment of genes associated with both presynaptic and postsynaptic structures amongst those contained within all four modules of interest (namely the substantia nigra “cyan”, putamen “cyan”, frontal cortex “lightyellow” and white matter “blue” modules). However, whereas the substantia nigra “cyan” and white matter “blue” modules were more significantly enriched for genes associated with presynaptic structures, the putamen “cyan” and frontal cortex “lightyellow” modules showed more significant enrichments for genes associated with postsynaptic structures (Fig. 3; **Supplementary Table 3**).

**Figure 3.**
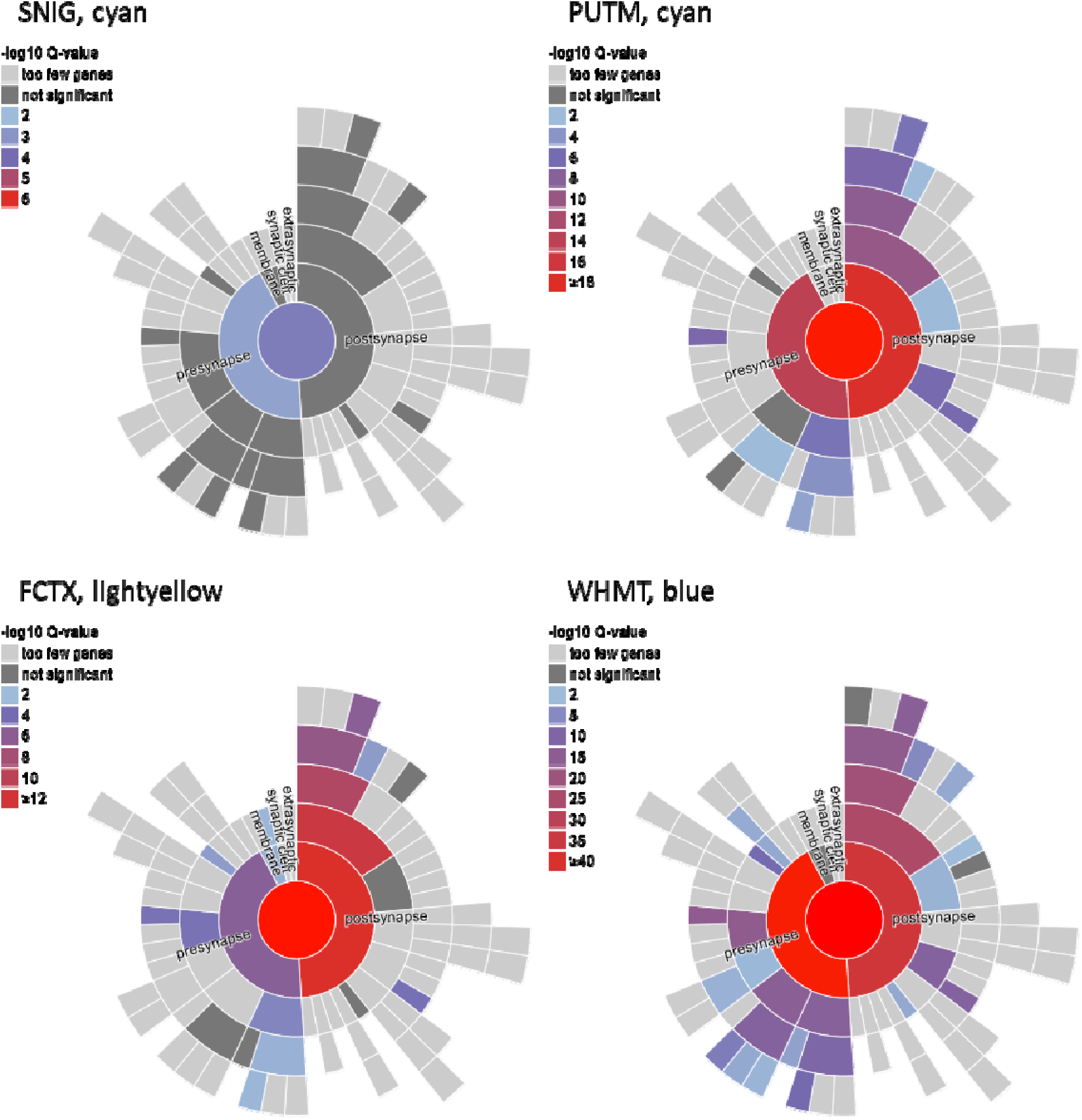
Variable enrichment of genes associated with presynaptic and postsynaptic structures within dystonia-linked UKBEC co-expression modules. Visualisations of the enrichment of SynGO ontology terms for synaptic location within the dystonia-linked UKBEC modules (the substantia nigra “cyan”, putamen “cyan”, frontal cortex “lightyellow” and white matter “blue” modules) provided by the SynGO web resource (https://www.syngoportal.org/index.html). The hierarchical structure of SynGO terms is represented by concentric rings with the most specific terms placed peripherally. The colour coding of the terms is based on the enrichment q-values. SNIG, substantia nigra; PUTM, putamen; FCTX, frontal cortex; WHMT, white matter.

### Dystonia-linked modules in the putamen and substantia nigra represent expression signatures which are unique to these tissues

We noted high overlaps amongst the DYT genes that enrich in putamen, frontal cortex and white matter co-expression modules, with three DYT genes appearing in all three of the relevant modules (*ANO3, HPCA* and *KCTD17*), and a further four DYT genes appearing in at least two of these modules (*KCNA1, CACNA1A, PRRT2*, and *SCN8A*). As expected, given their well-defined role in nigral dopamine synthesis, the genes clustering in the substantia nigra module were found to enrich exclusively in that tissue. We extended this analysis by calculating module preservation statistics to formally assess how similar the four dystonia-linked modules were to all co-expression modules identified in the UKBEC dataset. Interestingly, we found that while the white matter “blue” and frontal cortex “lightyellow” modules showed strong evidence of preservation in other brain regions, the substantia nigra and putamen “cyan” modules showed weak evidence of preservation (white matter “blue” module, median Z summary statistic = 56.71; frontal cortex “lightyellow” module, median Z summary statistic = 12.35; putamen “cyan” module, mean Z summary statistic = 5.93; substantia nigra “cyan” module, median Z summary statistic = 3.95) (Fig. 4**, Supplementary Table 4**). This indicates that the dystonia-linked putamen and substantia nigra co-expression modules represent expression signatures specific to these brain tissues. Furthermore, these results suggest that while the DYT genes contained within these modules may be widely expressed, they have gene-gene interactions which are tissue-specific in nature.

**Figure 4.**
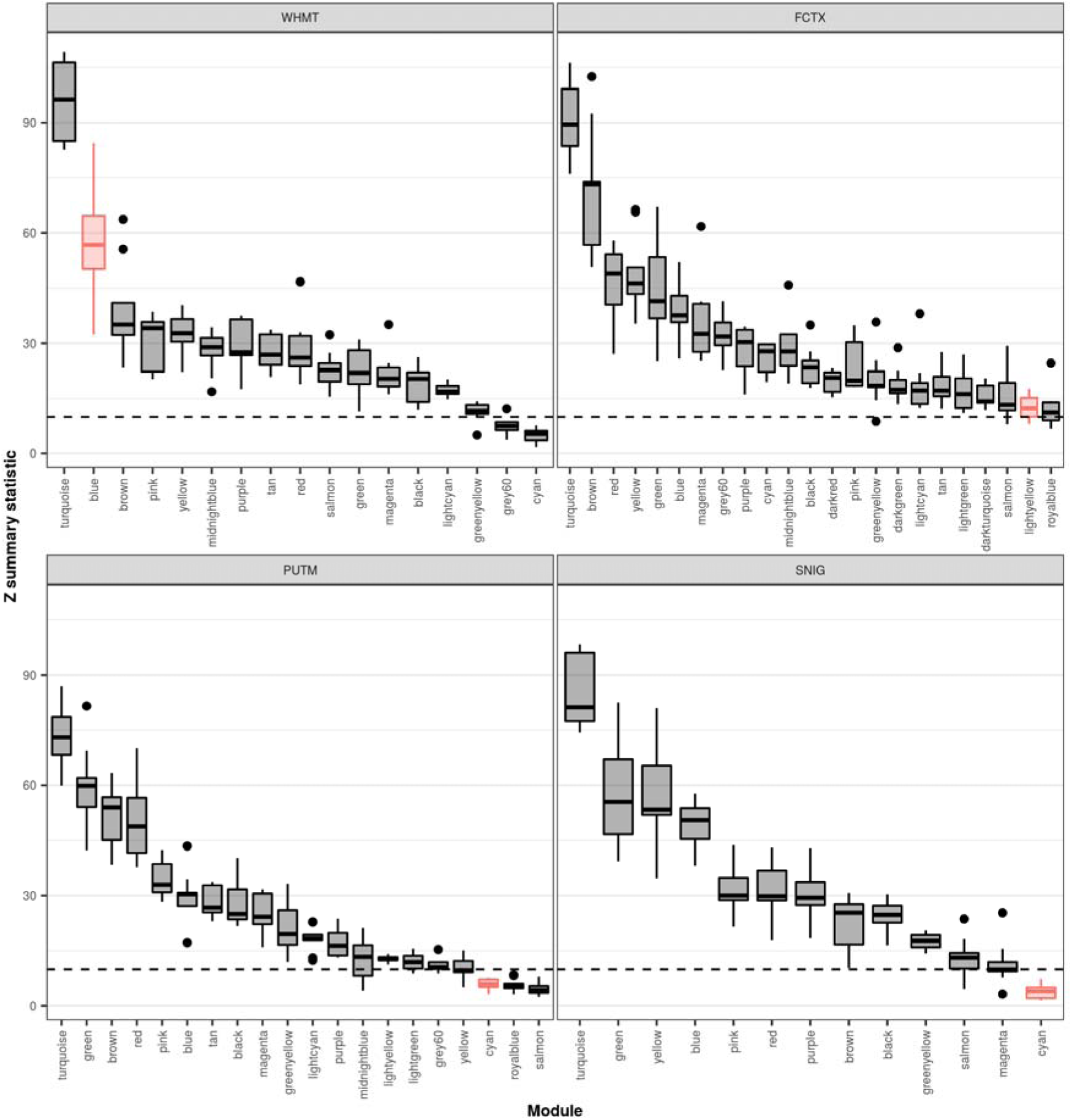
Module preservation within UKBEC tissues containing dystonia-linked modules. Plot of module preservation for each module within UKBEC tissues containing dystonia-linked modules. Module preservation, as denoted by the Z summary statistic, was determined using a WGCNA-based preservation analysis applied across all ten UKBEC co-expression networks. Thus, each boxplot represents the preservation of the named module within the labelled tissue across the remaining nine tissues. The black dashed lines indicate the threshold for strong evidence of module preservation (Z summary statistic > 10), as defined by Langfelder et al. (Langfelder *et al*., 2011). Modules enriched with dystonia genes are highlighted in red.

### Heritability of major-depressive disorder, obsessive-compulsive disorder and schizophrenia is significantly enriched in dystonia-linked modules

The identification of four UKBEC-derived co-expression modules enriched for DYT genes (substantia nigra “cyan”, putamen “cyan”, frontal cortex “lightyellow” and white matter “blue”) provided an opportunity to investigate commonalities in the underlying genetic architecture of dystonia and a range of neuropsychiatric disorders noted to have a high prevalence amongst individuals with the condition. Using stratified LDSC we tested whether genes assigned to the modules with high confidence (based on a module membership of > 0.5) significantly contributed to the common SNP heritability of anxiety, major-depressive disorder, obsessive-compulsive disorder, and schizophrenia. Given that in recent years dystonia has been viewed primarily as a movement disorder with significant phenotypic overlap with Parkinson disease (Shetty *et al*., 2019), we extended our analysis to include this condition.

Stratified LDSC demonstrated significant enrichments of obsessive-compulsive disorder and schizophrenia heritability in the putamen “cyan” (obsessive-compulsive disorder, co-efficient p-value = 0.0019; schizophrenia, co-efficient p-value = 0.000047) and a significant enrichment of major-depressive disorder and schizophrenia heritability in the white matter “blue” module (major-depressive disorder, co-efficient p-value = 0.00039; schizophrenia, co-efficient p-value = 9.14 x 10^-10^; Fig. 5**, Supplementary Table 5**). While no significant enrichment in heritability was observed for genes contained within the substantia nigra “cyan” module for any of the diseases we investigated, including Parkinson disease, the frontal cortex “lightyellow” module was nominally enriched for the heritability of major depressive disorder (co-efficient p-value = 0.018).

**Figure 5.**
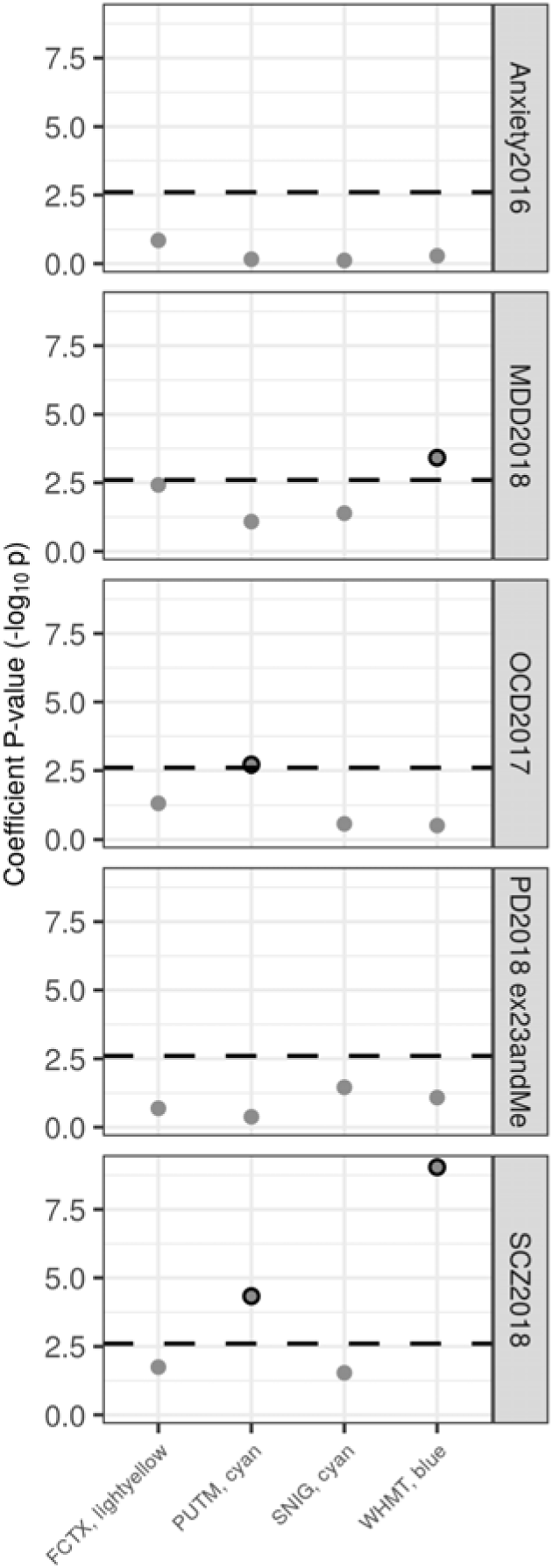
Enrichment of disease heritability within dystonia-linked UKBEC co-expression modules. Stratified LDSC using UKBEC co-expression modules. The black dashed lines indicate the cut-off for Bonferroni significance (*p* < 0.05/(4 x 5)). Bonferroni-significant results are marked with black borders. Numerical results are reported in **Supplementary Table 3**. FCTX, frontal cortex; PUTM, putamen; MDD, major depressive disorder; OCD, obsessive compulsive disorder; PD, Parkinson’s disease; SCZ, schizophrenia; SNIG, substantia nigra; WHMT, white matter.

## Discussion

While there has been remarkable progress in our understanding of the genetic structure of dystonia, the anatomical, cellular, and molecular basis remains unknown for most genetic forms of dystonia, as does its genetic and biological relationship to neuropsychiatric disorders. Using a systems biology approach leveraging our current understanding of the genetic basis of dystonia and neuropsychiatric disease, we show that: (i) the expression of the currently known DYT genes is significantly enriched in adult nigral dopaminergic neurons and striatal MSNs; (ii) multiple DYT genes are highly co-expressed with each other in multiple brain regions relevant to dystonia functional neuroanatomy, including the substantia nigra, putamen, frontal cortex and white matter; (iii) and finally, there is evidence of a genetic relationship between dystonia and complex neuropsychiatric diseases.

We found that the enrichment of DYT genes in adult nigral dopaminergic neurons detected using EWCE analysis was largely driven by genes that are involved in various steps of dopamine synthesis and metabolism in nigral dopaminergic neurons and when mutated are responsible for DOPA-responsive dystonias (Ribot *et al*., 2019). Similarly, WGCNA-based analyses showed that the same genes clustered together in a single co-expression module specific to the substantia nigra (“cyan” module). These results are consistent with expectation, thus highlighting the reliability of our hypothesis-free approach in identifying disease-relevant cell types and tissue-specific gene-gene interactions amongst DYT genes. With this in mind, we note that none of the “non-DOPA-responsive” DYT genes were highly specific to dopaminergic neurons, nor did we find that any co-clustered with the DOPA-responsive DYT genes in the substantia nigra “cyan” co-expression module. Together, these results confirm the growing view that DOPA-responsive dystonias should be considered a distinct subgroup of dystonia from both a clinical and biological perspective. Furthermore, our data indicate that non-DOPA-responsive DYT genes are unlikely to contribute to dystonia through dysfunction of dopamine metabolism in dopaminergic neurons.

Other neuronal cell types highlighted by our EWCE-and WGCNA-based analyses include MSNs, which constitute 95% of the cellular population of the putamen (Gerfen and Surmeier, 2011). EWCE analysis showed that four DYT genes, namely *ADCY5, GNAL, ANO3,* and *KCTD17,* had the highest specificity of expression in MSNs, suggesting they were the DYT genes driving the enrichment in MSNs. Importantly, WGCNA results went beyond simply suggesting a role for individual DYT genes within MSNs and indicated a functional interaction of multiple DYT genes in a single convergent pathway. Indeed, we observed eight DYT genes, including three out of the four DYT genes with the highest specificity in MSNs (*ADCY5, ANO3,* and *KCTD17*) and five others that were not highlighted by EWCE analysis (*CACNA1A, SCN8A, KCNA1, PRRT2, HPCA*), to be co-expressed in the putamen “cyan” module (the top module for DYT gene enrichment across all tested co-expression modules). This co-expression module also contained several established MSN-specific expression markers, including: *DRD1*, *DRD2* and *ADORA2A*, which encode the striatal dopamine and adenosine receptors; and *PPP1R1B*, which encodes DARPP-32, the universal marker of MSNs and part of the signalling cascade downstream of dopaminergic and adenosinergic receptor activation (Fienberg *et al*., 1998). In summary, these results support a functional interaction and biological convergence of multiple DYT genes in MSNs. This biological convergence is noteworthy, given that the function of several DYT genes found clustering in the putamen “cyan” co-expression module (e.g. *KCTD17*, *HPCA*, *ANO3*) is poorly characterized, especially in the context of MSN biology.

To better understand the biological function of DYT genes, all dystonia-linked co-expression modules were functionally annotated using Gene Ontology terms. This functional annotation revealed that all four dystonia-linked co-expression modules were enriched for genes associated with synaptic function (in particular, genes associated with presynaptic and postsynaptic structures), suggesting that DYT-genes found within these modules are involved too in synaptic function. Notably, in the putamen-specific “cyan” co-expression module the enrichment for genes associated with postsynaptic structures yield a lower p-value than that for genes associated with presynaptic structures. Given the central role of MSNs in receiving and gating synaptic inputs from cortical and thalamic glutamatergic neurons, this suggests that disruption of postsynaptic function in MSNs through mutations in multiple DYT genes may be important in dystonia pathogenesis. In support of this hypothesis are the following observations: (i) abnormal plasticity and loss of synaptic downscaling at cortico-striatal synapses has been shown to be a dystonia endophenotype shared by different genetic animal models of dystonia (Martella *et al*., 2014; Calabresi *et al*., 2016; Maltese *et al*., 2017; Zakirova *et al*., 2018; Yu-Taeger *et al*., 2019); (ii) *ADCY5* and *GNAL*, two DYT genes, form part of the signalling transduction machinery in response to stimulation of dopaminergic and adenosinergic signaling in MSNs (Herve, 2011; Goodchild *et al*., 2013); (iii) and finally, *Insomniac* and *nca* (Drosophila homologues of *KCTD17* and *HPCA,* respectively, both of which are DYT genes in the putamen “cyan” module) have been found to modulate sleep in *Drosophila* through disruption of dopaminergic post-synaptic signalling (Pfeiffenberger and Allada, 2012; Chen *et al*., 2019; Kikuma *et al*., 2019).

These findings have significant clinical implications, particularly for classification of dystonias. Currently, genetic forms of dystonia are primarily classified based on their clinical presentation. However, there is significant variability and pleiotropy in the clinical presentation of DYT mutation carriers (Table 1), which makes the exact assignment of a DYT mutation to a specific class of dystonia a hard task. Additionally, it is debatable whether grouping patients based on their clinical presentation correctly mirrors the underlying neuroanatomical or biological substrates of different types of dystonia. Importantly, the phenotypes associated with mutations in the DYT genes enriched in the putamen “cyan” module belonged to different clinical subtypes of dystonia, namely isolated dystonia (*ANO3* and *HPCA*), combined dystonia (*KCTD17* and *ADCY5*), and paroxysmal dystonia (*KCNA1, CACNA1A, PRRT2* and *SCN8A*). Furthermore, the putamen “cyan” module contained several other genes linked to monogenic hyperkinetic movement disorders, including genes associated with inherited forms of chorea (*PDE2A, GPR88, HTT, VPS13A, JPH3*) and for dyskinetic epileptic encephalopathies (*GNB1*, *UNC13A*, *GRIN1*, *STX1B*, *KNCQ2*, and *CACNA1B*), strongly suggesting that similar neuroanatomical and biological substrates underlie different clinical subtypes of monogenic dystonias and other hyperkinetic movement disorders. This finding is highly consistent with the growing appreciation that many neurogenetic disorders are characterised by genetic pleiotropy and variable expressivity (Warman Chardon *et al*., 2015), and indicates that the current system of dystonia classification based on clinical presentation may not reflect the molecular structure of the disease.

Clinically, variable expressivity may also extend to the neuropsychiatric symptoms often observed in individuals with dystonia. In support of this, we showed that the co-expression modules enriched for DYT genes also enriched for the heritability of several neuropsychiatric disorders, including major-depressive disorder, obsessive-compulsive disorder and schizophrenia. These results also reinforce the concept that neuropsychiatric disorders commonly observed in dystonic patients, such as major-depressive disorder and obsessive-compulsive disorder, are intrinsic to the neurobiology of dystonia. More specifically, these findings suggest that psychiatric symptoms are not merely a reaction to the disability arising from dystonia, but rather, the underlying molecular pathophysiology of dystonia increases a patient’s risk of developing psychiatric symptoms.

Notably, integration of GWAS-identified risk variants for obsessive-compulsive disorder and schizophrenia together with recent transcriptomic analyses have implicated MSNs in the neurobiology of these neuropsychiatric diseases (Skene *et al*., 2018; Yilmaz *et al*., 2018). We too observed an enrichment of obsessive-compulsive disorder and schizophrenia heritability in the MSN-related “cyan” putamen module, suggesting that dysfunction of MSN synaptic activity, resulting from different types of genetic insult, may represent an overlap in the biology of dystonia and these neuropsychiatric conditions. Similarly, glutamatergic pyramidal neurons have also been shown to be central to the aetiology of schizophrenia, and the “blue” white-matter module, which enriched for markers of pyramidal neurons, also enriched for DYT genes and schizophrenia heritability (Skene *et al*., 2018). Conversely, the enrichment of major-depressive disorder heritability was observed only for the frontal cortex “lightyellow” and “blue” white matter modules but not in the “cyan” putamen module. Given the known role of frontal and prefrontal cortex circuits in the pathogenesis of major-depressive disorder and other mood disorders (Price and Drevets, 2012), these results suggest that synaptic dysfunction induced by dysregulation of dystonia genes and their interactors in these particular brain regions might underpin the high risk and occurrence of depression in dystonia patients. These findings appear to support the concept that the same genetic disruption could operate across multiple brain regions and produce different clinical effects (i.e. dystonia vs predisposition to psychiatric symptoms), depending on the tissue-specific gene-gene interactions present.

### Limitations and Conclusions

While our analysis highlights the contribution of certain brain regions, cell-types and biological functions in dystonia pathogenesis, we recognise its limitations. First and foremost, this study depends on the quality and completeness of the genetic data we use. As we can only analyse known DYT genes to identify cell types and co-expression modules of interest, our findings could change with the identification of additional disease-associated genes. This limitation might explain our failure to detect an enrichment of DYT genes in other dystonia-associated brain regions (e.g. cerebellum). Similarly, limitations in GWAS power (in particular, for anxiety and obsessive-compulsive disorder) limit our ability to assess enrichments in heritability within specific gene sets. Secondly, this study is limited by the availability of high quality, region and cell-specific gene expression data. Due to its completeness, we use mouse-derived cellular profiles to assess cell specificity of gene expression of the DYT genes, but we appreciate there may be species differences. In addition, the co-expression networks analysed are generated through the analysis of a set of adult-derived brain regions, limiting the spatial resolution, and more importantly, preventing the analysis of potentially critical developmental windows. Hence, we are unable to analyse DYT genes which are likely to generate dystonia through a fundamental role during brain development, such as *TOR1A*, *THAP1* and *KMT2B* (Vasudevan *et al*., 2006; Zhao *et al*., 2013; Faundes *et al*., 2018). Finally, given that we do not have access to significant quantities of brain transcriptomic data from individuals with genetic forms of dystonia, we assume that DYT genes operate through a loss of their normal function rather than through novel gains of function.

In conclusion, this work enabled the unbiased identification of brain region-specific modules of biologically related genes and provided insights on cell-specific molecular signatures relevant to dystonia. We find that multiple DYT genes are functionally related in the adult human brain and likely contribute to modulation of synaptic signalling in striatal MSNs, adult dopaminergic neurons and frontal cortical neurons. While the exact mechanism of each individual DYT gene’s participation in this physiological process remains unknown, these results demonstrate a functional convergence of DYT genes linked to different phenotypic presentations and apparently unrelated cellular processes. These results bear significance for the treatment of dystonia, as future therapeutic approaches may target shared pathophysiological abnormalities, as opposed to symptoms. Finally, we demonstrate a genetic relationship between dystonia and several neuropsychiatric disorders, suggesting that disruption of genetic networks linked to dystonia pathogenesis in discrete brain regions may represent the neurobiological basis for the phenotypic overlap between dystonia and neuropsychiatric disorders.

## Appendix

### UK Brain Expression Consortium (UKBEC) members and affiliations

Juan A Botía (Department of Molecular Neuroscience, University College London (UCL) Institute of Neurology, London, UK; Department of Information and Communications Engineering, University of Murcia, Spain), Karishma D’Sa (Department of Molecular Neuroscience, University College London (UCL) Institute of Neurology, London, UK; Department of Medical & Molecular Genetics, King’s College London, Guy’s Hospital, London, UK), Paola Forabosco (Istituto di Ricerca Genetica e Biomedica, Cittadella Universitaria di Cagliari, 09042, Monserrato, Sardinia, Italy), Sebastian Guelfi (Department of Molecular Neuroscience, University College London (UCL) Institute of Neurology, London, UK), John Hardy (Department of Molecular Neuroscience, University College London (UCL) Institute of Neurology, London, UK), Jana Vandrovcova (Department of Molecular Neuroscience, University College London (UCL) Institute of Neurology, London, UK), Chris-Ann Mackenzie (Department of Neuropathology, MRC Sudden Death Brain Bank Project, University of Edinburgh, Edinburgh, UK), Adaikalavan Ramasamy (Department of Molecular Neuroscience, University College London (UCL) Institute of Neurology, London, UK; Jenner Institute, University of Oxford, Oxford, UK), Mina Ryten (Department of Molecular Neuroscience, University College London (UCL) Institute of Neurology, London, UK; Department of Medical & Molecular Genetics, King’s College London, Guy’s Hospital, London, UK), Colin Smith (Department of Neuropathology, MRC Sudden Death Brain Bank Project, University of Edinburgh, Edinburgh, UK), Daniah Trabzuni (Department of Molecular Neuroscience, University College London (UCL) Institute of Neurology, London, UK; Department of Genetics, King Faisal Specialist Hospital and Research Centre, Riyadh, Saudi Arabia), Michael E Weale (Department of Medical & Molecular Genetics, King’s College London, Guy’s Hospital, London, UK)

### IPDGC consortium members and affiliations

**<u>United Kingdom</u>:** <u>Alastair J Noyce</u> (Preventive Neurology Unit, Wolfson Institute of Preventive Medicine, QMUL, London, UK and Department of Molecular Neuroscience, UCL, London, UK), <u>Rauan Kaiyrzhanov</u> MD (Department of Molecular Neuroscience, UCL Institute of Neurology, London, UK), <u>Ben Middlehurst</u> (Institute of Translational Medicine, University of Liverpool, Liverpool, UK), <u>Demis A Kia</u> BSc (UCL Genetics Institute; and Department of Molecular Neuroscience, UCL Institute of Neurology, London, UK), <u>Manuela Tan</u> BA (Psych) (Department of Clinical Neuroscience, University College London, London, UK), <u>Henry Houlden</u> MD (Department of Molecular Neuroscience, UCL Institute of Neurology, London, UK), <u>Huw R Morris</u> PhD FRCP (Department of Clinical Neuroscience, University College London, London, UK), <u>Helene Plun-Favreau</u> PhD (Department of Molecular Neuroscience, UCL Institute of Neurology, London, UK), <u>Peter Holmans</u> PhD (Biostatistics & Bioinformatics Unit, Institute of Psychological Medicine and Clinical Neuroscience, MRC Centre for Neuropsychiatric Genetics & Genomics, Cardiff, UK), <u>John Hardy</u> PhD (Department of Molecular Neuroscience, UCL Institute of Neurology, London, UK), <u>Daniah Trabzuni</u> PhD (Department of Molecular Neuroscience, UCL Institute of Neurology, London, UK; Department of Genetics, King Faisal Specialist Hospital and Research Centre, Riyadh, 11211 Saudi Arabia), <u>Jose Bras</u> PhD (UK Dementia Research Institute at UCL and Department of Molecular Neuroscience, UCL Institute of Neurology, London, UK), <u>John Quinn</u> PhD (Institute of Translational Medicine, University of Liverpool, Liverpool, UK), <u>Kin Y Mok</u> FRCP PhD (Department of Molecular Neuroscience, UCL Institute of Neurology, London, UK; Division of Life Science, Hong Kong University of Science and Technology, Hong Kong SAR, China), <u>Kerri J. Kinghorn MBBChir PhD MRCP</u> (Institute of Healthy Ageing, University College London, London, UK), <u>Kimberley Billingsley</u> (Institute of Translational Medicine, University of Liverpool, Liverpool, UK), <u>Nicholas W Wood</u> PhD FRCP (UCL Genetics Institute; and Department of Molecular Neuroscience, UCL Institute of Neurology, London, UK), <u>Patrick Lewis</u> PhD (University of Reading, Reading, UK), <u>Rita Guerreiro</u> PhD (UK Dementia Research Institute at UCL and Department of Molecular Neuroscience, UCL Institute of Neurology, London, UK), <u>Ruth Lovering</u> PhD (University College London, London, UK), <u>Lea R’Bibo</u> (Department of Molecular Neuroscience, UCL Institute of Neurology, London, UK), <u>Claudia Manzoni PhD</u> (University of Reading, Reading, UK), <u>Mie Rizig</u> PhD (Department of Molecular Neuroscience, UCL Institute of Neurology, London, UK), <u>Mina Ryten</u> (MBBS PhD MRCP, Department of Molecular Neuroscience, UCL Institute of Neurology, London, UK), <u>Sebastian Guelfi</u> (BSc, Department of Molecular Neuroscience, UCL Institute of Neurology, London, UK), <u>Valentina Escott-Price</u> PhD (MRC Centre for Neuropsychiatric Genetics and Genomics, Dementia Research Institute, Cardiff University School of Medicine, Cardiff, UK), <u>Viorica Chelban</u> (Department of Molecular Neuroscience, UCL Institute of Neurology, London, UK), <u>Thomas Foltynie</u> (UCL Institute of Neurology, London, UK) MRCP PhD, <u>Nigel Williams</u> PhD (MRC Centre for Neuropsychiatric Genetics and Genomics, Cardiff, UK), Chingiz Shashakin PhD (Department of Molecular Neuroscience, Institute of Neurology, UCL, London, UK), Nazira Zharkinbekova (Department of Molecular Neuroscience, Institute of Neurology, UCL, London, UK) Elena Zholdybayeva PhD (Department of Molecular Neuroscience, Institute of Neurology, UCL, London, UK), Akbota Aitkulova PhD (Department of Molecular Neuroscience, Institute of Neurology, UCL, London, UK), <u>Kirsten Harvey</u> MD PhD (UCL School of Pharmacy, UK).

**<u>France:</u>** <u>Alexis Brice MD</u> (Institut du Cerveau et de la Moelle épinière, ICM, Inserm U 1127, CNRS, UMR 7225, Sorbonne Universités, UPMC University Paris 06, UMR S 1127, AP-HP, Pitié-Salpêtrière Hospital, Paris, France), <u>Fabrice Danjou</u> MD, PhD (Institut du Cerveau et de la Moelle épinière, ICM, Inserm U 1127, CNRS, UMR 7225, Sorbonne Universités, UPMC University Paris 06, UMR S 1127, AP-HP, Pitié-Salpêtrière Hospital, Paris, France), <u>Suzanne Lesage</u> PhD (Institut du Cerveau et de la Moelle épinière, ICM, Inserm U 1127, CNRS, UMR 7225, Sorbonne Universités, UPMC University Paris 06, UMR S 1127, AP-HP, Pitié-Salpêtrière Hospital, Paris, France), <u>Jean-Christophe Corvol MD, PhD</u> (Institut du Cerveau et de la Moelle épinière, ICM, Inserm U 1127, CNRS, UMR 7225, Sorbonne Universités, UPMC University Paris 06, UMR S 1127, Centre d’Investigation Clinique Pitié Neurosciences CIC-1422, AP-HP, Pitié-Salpêtrière Hospital, Paris, France), <u>Maria Martinez</u> PhD (INSERM UMR 1220; and Paul Sabatier University, Toulouse, France),

**<u>Germany</u>:** <u>Anamika Giri</u> M.Sc. (Department for Neurodegenerative Diseases, Hertie Institute for Clinical Brain Research, University of Tübingen, and DZNE, German Center for Neurodegenerative Diseases, Tübingen, Germany), <u>Claudia Schulte</u> (Department for Neurodegenerative Diseases, Hertie Institute for Clinical Brain Research, University of Tübingen, and DZNE, German Center for Neurodegenerative Diseases, Tübingen, Germany), <u>Kathrin Brockmann MD</u> (Department for Neurodegenerative Diseases, Hertie Institute for Clinical Brain Research, University of Tübingen, and DZNE, German Center for Neurodegenerative Diseases, Tübingen, Germany), <u>Javier Simón-Sánchez PhD</u> (Department for Neurodegenerative Diseases, Hertie Institute for Clinical Brain Research, University of Tübingen, and DZNE, German Center for Neurodegenerative Diseases, Tübingen, Germany), <u>Peter Heutink</u> PhD (DZNE, German Center for Neurodegenerative Diseases and Department for Neurodegenerative Diseases, Hertie Institute for Clinical Brain Research, University of Tübingen, Tübingen, Germany), <u>Patrizia Rizzu</u> PhD (DZNE, German Center for Neurodegenerative Diseases), <u>Manu Sharma</u> PhD (Centre for Genetic Epidemiology, Institute for Clinical Epidemiology and Applied Biometry, University of Tubingen, Germany), <u>Thomas Gasser</u> MD(Department for Neurodegenerative Diseases, Hertie Institute for Clinical Brain Research, and DZNE, German Center for Neurodegenerative Diseases, Tübingen, Germany),

**<u>United States of America:</u>** <u>Aude Nicolas</u> (Laboratory of Neurogenetics, National Institute on Aging, Bethesda, MD, USA), <u>Mark R Cookson</u> PhD (Laboratory of Neurogenetics, National Institute on Aging, Bethesda, USA), <u>Sara Bandres-Ciga</u> PhD (Laboratory of Neurogenetics, National Institute on Aging, Bethesda, MD, USA), <u>Cornelis Blauwendraat</u> PhD (National Institute on Aging and National Institute of Neurological Disorders and Stroke, USA), <u>David W. Craig</u> (Department of Translational Genomics, Keck School of Medicine, University of Southern California, Los Angeles, USA), <u>Faraz Faghri</u> MS (Laboratory of Neurogenetics, National Institute on Aging, Bethesda, USA; Department of Computer Science, University of Illinois at Urbana-Champaign, Urbana, IL, USA), <u>J Raphael Gibbs</u> PhD (Laboratory of Neurogenetics, National Institute on Aging, National Institutes of Health, Bethesda, MD, USA), <u>Dena G. Hernandez</u> PhD (Laboratory of Neurogenetics, National Institute on Aging, Bethesda, MD, USA), <u>Kendall Van Keuren-Jensen</u> (Neurogenomics Division, TGen, Phoenix, AZ USA), <u>Joshua M. Shulman MD, PhD</u> (Departments of Neurology, Neuroscience, and Molecular & Human Genetics, Baylor College of Medicine, Houston, Texas, USA; Jan and Dan Duncan Neurological Research Institute, Texas Children’s Hospital, Houston, Texas, USA), <u>Hampton L. Leonard</u> (Laboratory of Neurogenetics, National Institute on Aging, Bethesda, MD, USA), <u>Mike A. Nalls</u> PhD (Laboratory of Neurogenetics, National Institute on Aging, Bethesda, USA; CEO/Consultant Data Tecnica International, Glen Echo, MD, USA), <u>Laurie Robak MD, PhD</u> (Baylor College of Medicine, Houston, Texas, USA), <u>Steven Lubbe PhD</u> (Ken and Ruth Davee Department of Neurology, Northwestern University Feinberg School of Medicine, Chicago, IL, USA), <u>Steven Finkbeiner</u> (Departments of Neurology and Physiology, University of California, San Francisco; Gladstone Institute of Neurological Disease; Taube/Koret Center for Neurodegenerative Disease Research, San Francisco, CA, USA), <u>Niccolo E. Mencacci</u> MD, PhD (Northwestern University Feinberg School of Medicine, Chicago, IL, USA), <u>Codrin Lungu</u> (National Institutes of Health Division of Clinical Research, NINDS, National Institutes of Health, Bethesda, MD, USA), <u>Andrew B Singleton</u> PhD (Laboratory of Neurogenetics, National Institute on Aging, Bethesda, MD, USA), <u>Sonja W. Scholz</u> MD, PhD (Neurodegenerative Diseases Research Unit, National Institute of Neurological Disorders and Stroke, Bethesda, MD, USA), <u>Xylena Reed</u> PhD (Laboratory of Neurogenetics, National Institute on Aging, Bethesda, MD, USA). <u>Kendall Van Keuren-Jensen</u> (Neurogenomics Division, TGen, Phoenix, AZ, USA).

**<u>Canada</u>:** <u>Ziv Gan-Or</u> MD, PhD, (Montreal Neurological Institute and Hospital, Department of Neurology & Neurosurgery, Department of Human Genetics, McGill University, Montréal, QC, H3A 0G4, Canada), <u>Guy A. Rouleau</u> MD, PhD, (Montreal Neurological Institute and Hospital, Department of Neurology & Neurosurgery, Department of Human Genetics, McGill University, Montréal, QC, H3A 0G4, Canada)

**<u>The Netherlands</u>:** <u>Jacobus J van Hilten</u> (Department of Neurology, Leiden University Medical Center, Leiden, Netherlands), <u>Johan Marinus</u> (Department of Neurology, Leiden University Medical Center, Leiden, Netherlands)

**<u>Spain</u>:** <u>Astrid D. Adarmes-Gómez</u> (Instituto de Biomedicina de Sevilla (IBiS), Hospital Universitario Virgen del Rocío/CSIC/Universidad de Sevilla, Seville), <u>Miquel Aguilar</u> (Fundació Docència i Recerca Mútua de Terrassa and Movement Disorders Unit, Department of Neurology, University Hospital Mutua de Terrassa, Terrassa, Barcelona.),<u>Ignacio Alvarez</u> (Fundació Docència i Recerca Mútua de Terrassa and Movement Disorders Unit, Department of Neurology, University Hospital Mutua de Terrassa, Terrassa, Barcelona.),<u>Victoria Alvarez</u> (Hospital Universitario Central de Asturias, Oviedo), <u>Francisco Javier Barrero</u> (Hospital Universitario Parque Tecnologico de la Salud, Granada), <u>Jesús Alberto Bergareche Yarza</u> (Instituto de Investigación Sanitaria Biodonostia, San Sebastián), <u>Inmaculada Bernal-Bernal</u> (Instituto de Biomedicina de Sevilla (IBiS), Hospital Universitario Virgen del Rocío/CSIC/Universidad de Sevilla, Seville), <u>Marta Blazquez</u> (Hospital Universitario Central de Asturias, Oviedo), <u>Marta Bonilla-Toribio</u> (Instituto de Biomedicina de Sevilla (IBiS), Hospital Universitario Virgen del Rocío/CSIC/Universidad de Sevilla, Seville), <u>Juan A. Botía PhD,</u> (Universidad de Murcia, Murcia), <u>María Teresa Boungiorno</u> (Fundació Docència i Recerca Mútua de Terrassa and Movement Disorders Unit, Department of Neurology, University Hospital Mutua de Terrassa, Terrassa, Barcelona.) <u>Dolores Buiza-Rueda</u> (Instituto de Biomedicina de Sevilla (IBiS), Hospital Universitario Virgen del Rocío/CSIC/Universidad de Sevilla, Seville), <u>Ana Cámara</u> (Hospital Clinic de Barcelona), <u>Maria Carcel</u> (Fundació Docència i Recerca Mútua de Terrassa, Barcelona and Department of Neurology, Movement Disorders Unit, Hospital Universitari Mutua de Terrassa, Barcelona), <u>Fátima Carrillo</u> (Instituto de Biomedicina de Sevilla (IBiS), Hospital Universitario Virgen del Rocío/CSIC/Universidad de Sevilla, Seville), <u>Mario Carrión-Claro</u> (Instituto de Biomedicina de Sevilla (IBiS), Hospital Universitario Virgen del Rocío/CSIC/Universidad de Sevilla, Seville), <u>Debora Cerdan</u> (Hospital General de Segovia, Segovia), <u>Jordi Clarimón</u> PhD, (Memory Unit, Department of Neurology, IIB Sant Pau, Hospital de la Santa Creu i Sant Pau, Universitat Autònoma de Barcelona and Centro de Investigación Biomédica en Red en Enfermedades Neurodegenerativas (CIBERNED), Madrid),<u>Yaroslau Compta</u> (Hospital Clinic de Barcelona), <u>Monica Diez-Fairen</u> (Fundació Docència i Recerca Mútua de Terrassa and Movement Disorders Unit, Department of Neurology, University Hospital Mutua de Terrassa, Terrassa, Barcelona.), <u>Oriol Dols-Icardo</u> PhD, (Memory Unit, Department of Neurology, IIB Sant Pau, Hospital de la Santa Creu i Sant Pau, Universitat Autònoma de Barcelona, Barcelona, and Centro de Investigación Biomédica en Red en Enfermedades Neurodegenerativas (CIBERNED), Madrid), <u>Jacinto Duarte</u> (Hospital General de Segovia, Segovia), <u>Raquel Duran</u> (Centro de Investigacion Biomedica, Universidad de Granada, Granada), <u>Francisco Escamilla-Sevilla</u> (Hospital Universitario Virgen de las Nieves, Instituto de Investigación Biosanitaria de Granada, Granada), <u>Mario Ezquerra</u> (Hospital Clinic de Barcelona),Manel Fernández (Hospital Clinic de Barcelona), <u>Rubén Fernández-Santiago</u> (Hospital Clinic de Barcelona), <u>Ciara Garcia</u> (Hospital Universitario Central de Asturias, Oviedo), <u>Pedro García-Ruiz</u> (Instituto de Investigación Sanitaria Fundación Jiménez Díaz, Madrid), <u>Pilar Gómez-Garre</u> (Instituto de Biomedicina de Sevilla (IBiS), Hospital Universitario Virgen del Rocío/CSIC/Universidad de Sevilla, Seville), <u>Maria Jose Gomez Heredia</u> (Hospital Universitario Virgen de la Victoria, Malaga), I<u>sabel Gonzalez-Aramburu</u> (Hospital Universitario Marqués de Valdecilla-IDIVAL, Santander), <u>Ana Gorostidi Pagola</u> (Instituto de Investigación Sanitaria Biodonostia, San Sebastián), <u>Janet Hoenicka</u> (Institut de Recerca Sant Joan de Déu, Barcelona), <u>Jon Infante</u> (Hospital Universitario Marqués de Valdecilla-IDIVAL and University of Cantabria, Santander, and Centro de Investigación Biomédica en Red en Enfermedades Neurodegenerativas (CIBERNED), <u>Silvia Jesús</u> (Instituto de Biomedicina de Sevilla (IBiS), Hospital Universitario Virgen del Rocío/CSIC/Universidad de Sevilla, Seville), <u>Adriano Jimenez-Escrig</u> (Hospital Universitario Ramón y Cajal, Madrid), <u>Jaime Kulisevsky</u> MD, PhD,(Movement Disorders Unit, Department of Neurology, IIB Sant Pau, Hospital de la Santa Creu i Sant Pau, Universitat Autònoma de Barcelona, Barcelona, and Centro de Investigación Biomédica en Red en Enfermedades Neurodegenerativas (CIBERNED)), <u>Miguel A. Labrador-Espinosa</u> (Instituto de Biomedicina de Sevilla (IBiS), Hospital Universitario Virgen del Rocío/CSIC/Universidad de Sevilla, Seville), <u>Jose Luis Lopez-Sendon</u> (Hospital Universitario Ramón y Cajal, Madrid), <u>Adolfo López de Munain Arregui</u> (Instituto de Investigación Sanitaria Biodonostia, San Sebastián), <u>Daniel Macias</u> (Instituto de Biomedicina de Sevilla (IBiS), Hospital Universitario Virgen del Rocío/CSIC/Universidad de Sevilla, Seville), <u>Juan Marín</u> MD, (Movement Disorders Unit, Department of Neurology, IIB Sant Pau, Hospital de la Santa Creu i Sant Pau, Universitat Autònoma de Barcelona, Barcelona, and Centro de Investigación Biomédica en Red en Enfermedades Neurodegenerativas (CIBERNED)), <u>Maria Jose Marti</u> (Hospital Clinic Barcelona), <u>Juan Carlos Martínez-Castrillo</u> (Instituto Ramón y Cajal de Investigación Sanitaria, Hospital Universitario Ramón y Cajal, Madrid), <u>Carlota Méndez-del-Barrio</u> (Instituto de Biomedicina de Sevilla (IBiS), Hospital Universitario Virgen del Rocío/CSIC/Universidad de Sevilla, Seville), <u>Manuel Menéndez González</u> (Hospital Universitario Central de Asturias, Oviedo), Adolfo Mínguez (Hospital Universitario Virgen de las Nieves, Granada, Instituto de Investigación Biosanitaria de Granada), <u>Pablo Mir</u> (Instituto de Biomedicina de Sevilla (IBiS), Hospital Universitario Virgen del Rocío/CSIC/Universidad de Sevilla, Seville), <u>Elisabet Mondragon Rezola</u> (Instituto de Investigación Sanitaria Biodonostia, San Sebastián), <u>Esteban Muñoz</u> (Hospital Clinic Barcelona), <u>Javier Pagonabarraga</u> MD, PhD,(Movement Disorders Unit, Department of Neurology, IIB Sant Pau, Hospital de la Santa Creu i Sant Pau, Universitat Autònoma de Barcelona, Barcelona, and Centro de Investigación Biomédica en Red en Enfermedades Neurodegenerativas (CIBERNED)), <u>Pau Pastor</u> (Fundació Docència i Recerca Mútua de Terrassa and Movement Disorders Unit, Department of Neurology, University Hospital Mutua de Terrassa, Terrassa, Barcelona.), <u>Francisco Perez Errazquin</u> (Hospital Universitario Virgen de la Victoria, Malaga), T<u>eresa Periñán-Tocino</u> (Instituto de Biomedicina de Sevilla (IBiS), Hospital Universitario Virgen del Rocío/CSIC/Universidad de Sevilla, Seville), <u>Javier Ruiz-Martínez</u> (Hospital Universitario Donostia, Instituto de Investigación Sanitaria Biodonostia, San Sebastián), <u>Clara Ruz</u> (Centro de Investigacion Biomedica, Universidad de Granada, Granada), <u>Antonio Sanchez Rodriguez</u> (Hospital Universitario Marqués de Valdecilla-IDIVAL, Santander), <u>María Sierra</u> (Hospital Universitario Marqués de Valdecilla-IDIVAL, Santander), <u>Esther Suarez-Sanmartin</u> (Hospital Universitario Central de Asturias, Oviedo), <u>Cesar Tabernero</u> (Hospital General de Segovia, Segovia), <u>Juan Pablo Tartari</u> (Fundació Docència i Recerca Mútua de Terrassa and Movement Disorders Unit, Department of Neurology, University Hospital Mutua de Terrassa, Terrassa, Barcelona<u>), Cristina Tejera-Parrado</u> (Instituto de Biomedicina de Sevilla (IBiS), Hospital Universitario Virgen del Rocío/CSIC/Universidad de Sevilla, Seville), <u>Eduard Tolosa</u> (Hospital Clinic Barcelona), <u>Francesc Valldeoriola</u> (Hospital Clinic Barcelona), <u>Laura Vargas-González</u> (Instituto de Biomedicina de Sevilla (IBiS), Hospital Universitario Virgen del Rocío/CSIC/Universidad de Sevilla, Seville), <u>Lydia Vela</u> (Hospital Universitario Fundación Alcorcón, Madrid), <u>Francisco Vives</u> (Centro de Investigacion Biomedica, Universidad de Granada, Granada).

**<u>Austria:</u>** <u>Alexander Zimprich</u> MD (Department of Neurology, Medical University of Vienna, Austria)

**<u>Norway</u>:** <u>Lasse Pihlstrom MD, PhD</u> (Department of Neurology, Oslo University Hospital, Oslo, Norway)

**<u>Estonia</u>**: <u>Pille Taba</u> MD, PhD, (Department of Neurology and Neurosurgery, University of Tartu, Tartu, Estonia)

**<u>Australia</u>:** <u>Sulev Koks</u> MD, PhD, (Centre for Molecular Medicine and Innovative Therapeutics, Murdoch University, Murdoch, 6150, Perth, Western Australia; The Perron Institute for Neurological and Translational Science, Nedlands, 6009, Perth, Western Australia)

## Abbreviations

MSNs: Medium-spiny neurons
EWCE: Expression-weighted cell-type enrichment
WGCNA: Weighted Gene Co-Expression Analysis
LDSC: linkage disequilibrium score regression

## Description of Supplementary Data

### Supplementary Tables

Supplementary Table 1. Cell-specific enrichments p-values of dystonia genes calculated using EWCE

Supplementary Table 2. Enrichment p-values of dystonia genes within gene co-expression modules generated using brain transcriptomic data from the UK Brain Expression Consortium and the GTEx consortium

Supplementary Table 3. Enrichment p-values of SynGO annotations within the substantia nigra “cyan”, putamen “cyan”, frontal cortex “lightyellow” and white matter “blue” modules.

Supplementary Table 4. Module preservation across four UKBEC tissues containing dystonia-linked modules.

Supplementary Table 5. Results of stratified LDSC analysis across all gene co-expression modules of interest and a range of neuropsychiatric and neurodegenerative disorders

### Supplementary Figures

**Supplementary Figure 1. Cell-specific profiles of dystonia-associated genes.** Plots of specificity values for all dystonia-associated genes within level 1 cell types from the Karolinska single-cell RNA-sequencing superset. Specificity values were derived from Skene et al.^2^ who calculated specificity by dividing the mean expression of a gene in one cell type by the mean expression in all cell types. In other words, specificity is the proportion of a gene’s total expression attributable to one cell type, with a value of 0 meaning a gene is not expressed in that cell type and a value of 1 meaning that a gene is only expressed in that cell type. For each gene, the top 3 cell types, as ranked by specificity, are marked with black borders.

