## Supplementary figures and images for "Transcriptomic analysis of dystonia-associated genes reveals functional convergence within specific cell types and shared neurobiology with psychiatric disorders"

### Supplemental figure 1

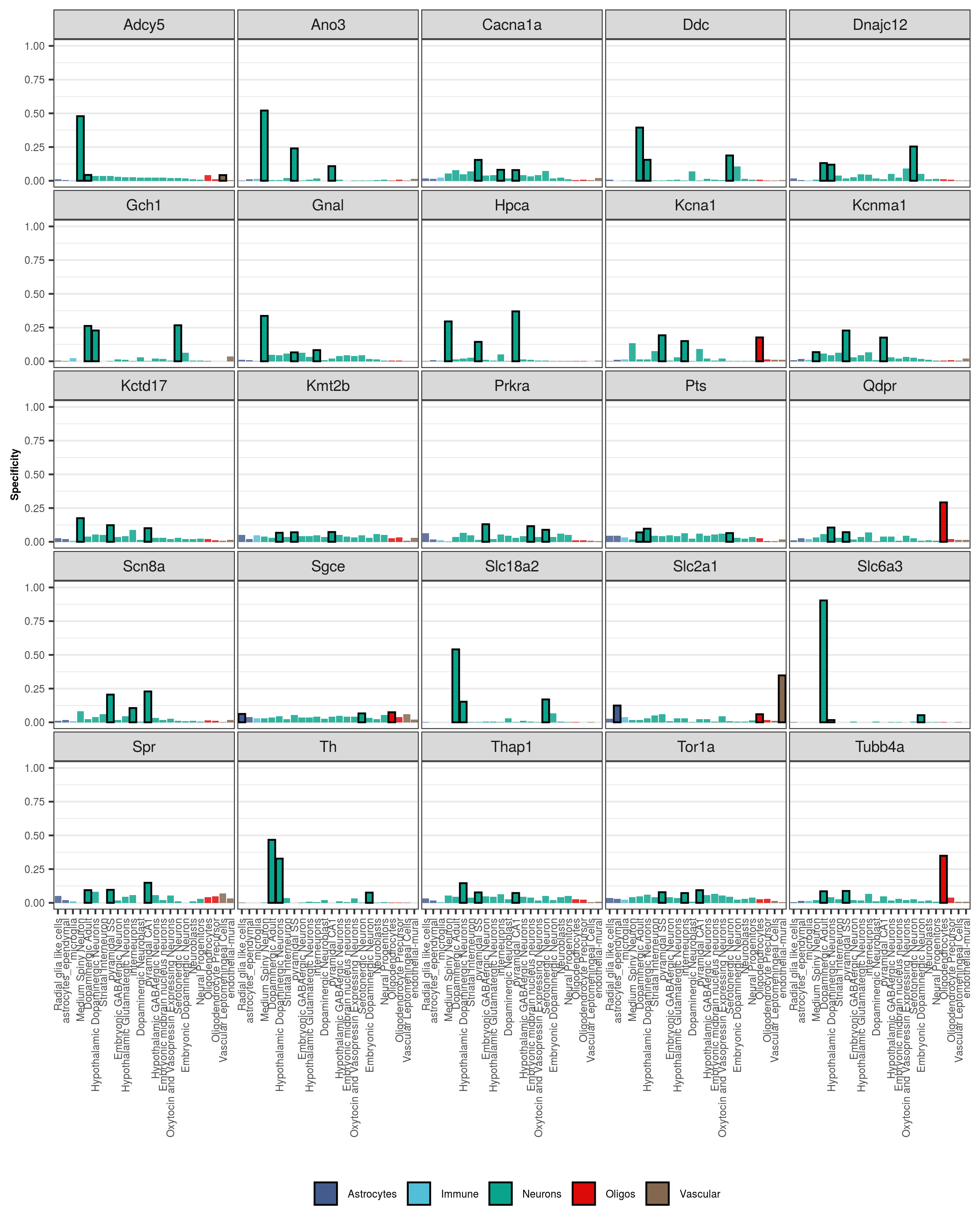
